# Individual Variation in Intrinsic Neuronal Properties of Nucleus Accumbens Core and Shell Medium Spiny Neurons in Animals Prone to Sign- or Goal-Track

**DOI:** 10.1101/2025.03.24.644332

**Authors:** Cristina E. María-Ríos, Geoffrey G. Murphy, Jonathan D. Morrow

## Abstract

The “sign-tracking” and “goal-tracking” model of individual variation in associative learning permits the identification of rats with different cue-reactivity and predisposition to addiction-like behaviors. Certainly, compared to “goal-trackers” (GTs), “sign-trackers” (STs) show more susceptibility traits such as increased cue-induced ‘relapse’ of drugs of abuse. Different cue- and reward-evoked patterns of activity in the nucleus accumbens (NAc) have been a hallmark of the ST/GT phenotype. However, it is unknown whether differences in the intrinsic neuronal properties of NAc medium spiny neurons (MSNs) in the core and shell subregions are also a physiological correlate of these phenotypes. We performed whole-cell slice electrophysiology in outbred male rats and found that STs exhibited the lowest excitability in the NAc core, with lower number of action potentials and firing frequency as well as a blunted voltage/current relationship curve in response to hyperpolarized potentials in both the NAc core and shell. Although firing properties of shell MSNs did not differ between STs and GTs, intermediate responders that engage in both behaviors showed greater excitability compared to both STs and GTs. These findings suggest that intrinsic excitability in the NAc may contribute to individual differences in the attribution of incentive salience.

**Significance Statement:** During associative learning, cues acquire predictive value, but in some instances, they also acquire incentive salience, meaning they take on some of the motivational properties of the reward. The propensity to attribute cues with incentive salience varies between individuals, and excessive attribution can lead to maladaptive behaviors. The “sign-and goal-tracking” model allows us to isolate these two properties and disambiguate the neurobiological processes that govern them. To our knowledge this is the first study characterizing passive and active membrane properties of MSNs in the NAc core and shell of STs and GTs, as well as IRs. These findings are meant to better inform investigations of the distinct role of the NAc in reward learning, particularly in the attribution of incentive salience and addiction predisposition.

## Introduction

The ability to develop adaptive responses to cues signaling food, safety, mating opportunities, and other rewards is essential for survival. The nucleus accumbens (NAc) plays a central role in this learning process. Individual variations in how information about the environment is processed through the NAc could contribute to traits associated with neuropsychiatric disorders including impulsivity, hyperactivity, and cue-reactivity (Robinson and Berridge, 1993; Berridge et al., 2009; Flagel and Robinson, 2017). However, linking such behaviorally complex predisposing factors to specific neurobiological processes has proven difficult. Because cues normally acquire both predictive and incentive value together, it is difficult to dissociate these properties and disambiguate the neurobiological processes that govern them. Such a dissociation can be achieved in rats by using Pavlovian conditioned approach (PavCA) procedures, which reveal individual differences in associative learning styles known as “sign-tracking” and “goal-tracking” (María-Ríos et al., 2023).

During Pavlovian conditioning, “sign-trackers” (STs) reliably approach the reward-paired cue and interact with it (“sign-tracking”), whereas “goal-trackers” (GTs) will direct their behavior away from the cue and towards the site of impending reward delivery (“goal-tracking”). The key difference between these phenotypes is that GTs only use reward-paired cues as predictors, but STs also attribute the cues with incentive salience, meaning the cues become irresistible and rewarding in and of themselves (Berridge et al., 2009). Intermediate responders (IRs) are rats that exhibit a relatively low bias toward sign- or goal-tracking behavior and tend to alternate between both conditioned responses. Compared to GTs, STs are more impulsive (Lovic et al., 2011), have less attentional control (Paolone et al., 2013), and are more susceptible to cue-induced “relapse” of drug self-administration (Saunders and Robinson, 2011; Versaggi et al., 2016) making it a useful model for studying the neurobiological basis of incentive salience attribution and its contribution to disorders like addiction.

Neuronal activity in the NAc seems to be particularly important for individual differences in incentive salience attribution. STs show increased cue-evoked c-fos expression in the NAc compared to GTs (Flagel et al., 2011a), and single-unit recordings and *in-vivo* voltammetry have also revealed that STs and GTs exhibit different cue- and reward-evoked patterns of activity and dopamine release in the NAc during Pavlovian learning (Flagel et al., 2011b; Gillis and Morrison, 2019). Much research in STs and GTs has focused on dissecting the molecular and neurochemical basis of these differences (for review see Flagel and Robinson, 2017). However, whether these variations in NAc activity reflect fundamental differences in the intrinsic neuronal excitability of NAc neurons remains unclear.

The NAc consists of two subregions known as the core and shell with distinct anatomical features (Záborszky et al., 1985; Zahm and Heimer, 1990; Heimer et al., 1991; Meredith et al., 1992; Britt et al., 2012) and functional roles in Pavlovian learning and motivation (Zahm, 1999; Bassareo et al., 2002; Day and Carelli, 2007; West and Carelli, 2016). GABAergic projection neurons known as medium spiny neurons (MSNs) comprise about 95% of the total neuronal population in both the core and the shell (Pennartz et al., 1994; Matamales et al., 2009). Therefore, the intrinsic excitability state of MSNs – how sensitive they are to changes in membrane potential caused by input stimuli – may reveal important functional differences between STs, GTs, and IRs in how the NAc integrates and relays reward information (O’Donnell and Grace, 1996; Nicola et al., 2000; Planert et al., 2013; Dorris et al., 2015).

The present study used whole-cell patch-clamp recordings to conduct an electrophysiological characterization of the passive and active membrane properties of MSNs in the NAc core and shell of ST, GT, and IR rats. We hypothesize that MSNs from these three phenotypes will express basal differences in intrinsic excitability and that these will contribute to our understanding of how the NAc, particularly of STs and GTs, differentially encodes predictive and incentive value of appetitive cues.

## Materials and methods

### Animals

Thirty-one adult male Sprague Dawley rats (aged 8-10 weeks) were purchased from Charles River Laboratories (C72, R04) and were housed in pairs. The rats followed a 12:12-hour light/dark cycle, with ad libitum access to food and water throughout the experiment. All rats were acclimatized to the housing colony for at least two days prior to handling. Following seven days of behavioral testing, the rats were allowed a resting period of 1-3 weeks in their home cages before electrophysiological recordings (**Figure 1**). All procedures involving animals had received prior approval from the University Committee on the Use and Care of Animals at the University of Michigan, Ann Arbor, MI.

**Figure 1.**
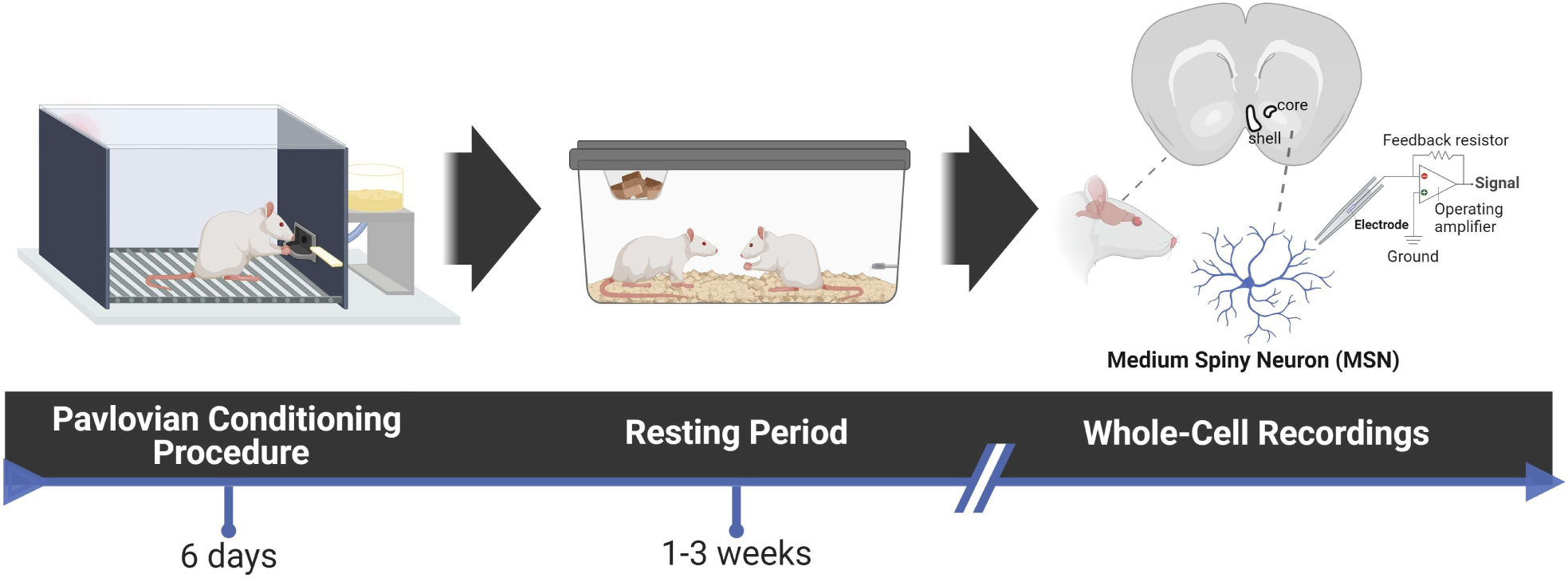
Experimental timeline. Rats underwent a 6-day Pavlovian conditioned approach (PavCA) procedure, where a neutral lever cue (CS) was paired with a banana-flavored pellet reward (US) following CS retraction. Each session included 25 CS-US pairings (ITI: 30-60 s). After PavCA, rats rested for 1-3 weeks before undergoing whole-cell recordings of medium spiny neurons (MSNs) in the nucleus accumbens core and shell.

### Drugs

Isoflurane (Fluriso - VetOne; Boise, ID) was administered at 5% via inhalation for inducing anesthesia. Unless otherwise stated, all chemicals were purchased from Tocris Bioscience (Bristol, UK), Sigma-Aldrich (St. Louis, MO), and Fisher Chemical (Pittsburgh, PA).

### Behavioral Testing Apparatus

As previously described (Maria-Rios et al., 2023), sixteen operant conditioning chambers with dimensions of 24.1 cm in width, 20.5 cm in depth, and 29.2 cm in height (manufactured by MED Associates, Inc.; St. Albans, VT) were utilized for behavioral testing. Each chamber was placed within a sound-attenuating cubicle equipped with a ventilation fan for ambient background noise during testing. Every chamber was equipped with a food magazine, a retractable lever (counterbalanced on either the left or right side of the magazine), and a red house light situated on the wall opposite to the magazine. Magazine entries were detected by an infrared sensor, and the levers were calibrated to register deflections in response to a minimum applied weight of 10 g. The inside of the lever mechanism featured a mounted LED to illuminate the slot through which the lever extended each time it protruded into the chamber. ABET II Software (Lafayette Instrument; Lafayette, IN) was used to automatically record the number and latency of lever presses and magazine entries.

### Behavioral Testing Procedure

Pavlovian conditioning was performed as previously described (María-Ríos et al., 2023). For two days prior to the start of training, all rats were habituated to the handler and the food reward. Rats were handled individually and were given banana-flavored pellets (45 mg; Bio-Serv; Frenchtown, NJ) in their home cages. On the third day, the rats were subjected to one pre-training session in the test chambers. Twenty-five food pellets were delivered on a variable time (VT) 30-s schedule (i.e., one pellet was delivered on average every 30 s, but varied 0-60 s). During this session, the lever remained retracted. Following pre-training, all rats underwent six daily sessions of behavioral training (**Figure 2A-C**). Each trial consisted of a presentation of the illuminated lever (conditioned stimulus, CS) into the chamber for 10 seconds on a VT 45-s schedule (i.e., time randomly varied 30-60 s between CS presentations). Immediately after retraction of the lever, there was a response-independent delivery of one pellet into the magazine (unconditioned stimulus, US). The beginning of the next inter-trial interval (ITI) began once both the lever and the pellet had been presented, and each test session consisted of 25 trials of CS-US pairings. All rats consumed all the pellets that were delivered. Rats were not food deprived at any point during experimentation.

**Figure 2.**
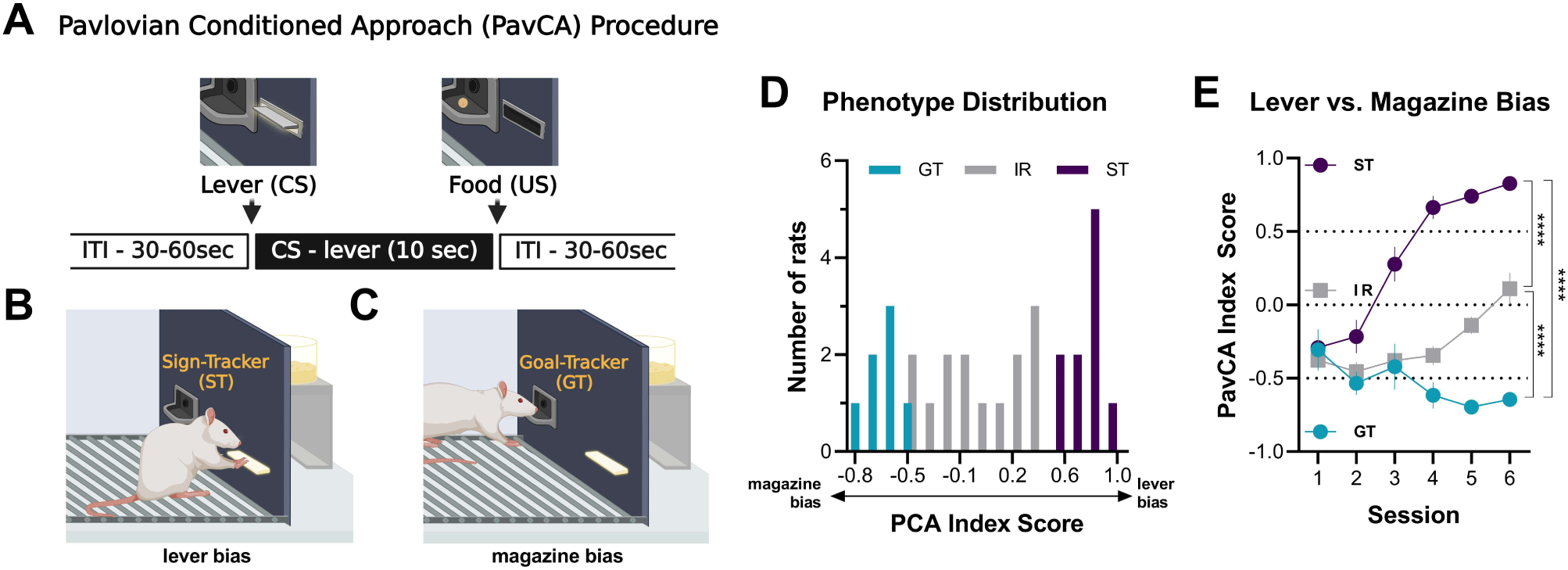
Classification of Pavlovian conditioned approach phenotypes. Rats were classified as sign-trackers (ST; n =10), goal-trackers (GT; n = 7), or intermediate responders (IR; n = 14) based on their lever and magazine bias during PavCA. (**A**) Each PavCA trial involved the extension of a lever-cue (CS) followed by food pellet delivery (US) after 10 seconds (inter-trial interval: 30-60 s). (**B**) STs approach the lever during the CS, despite no need for interaction for reward. (**C**) GTs approach the magazine, the site of food delivery, in response to the CS. (**D**) Distribution of ST (score ≥ 0.5), GT (score ≤ -0.5), and IR (-0.5 < score < 0.5) phenotypes. (**E**) PavCA index scores for ST, GT, and IR groups across six training sessions, showing significant differences between phenotypes. Data are presented as mean ± S.E.M. Significance for Sidak’s post hoc test is shown as **** p < 0.0001.

### Electrophysiology

#### Slice Preparation

After a 1-3-week resting period following the final day of PavCA training, slice preparation was performed as previously described (Maria-Rios et al., 2023). All rats were euthanized by decapitation after inducing anesthesia with isoflurane (Kent Scientific; Torrington, CT). Following rapid brain dissection, the brain was glued on a platform, which was then submerged in an ice-cold oxygenated (95% O2/ 5% CO2) cutting solution containing (in mM): 206 sucrose, 10 d-glucose, 1.25 NaH2PO4, 26 NaHCO3, 2 KCl, 0.4 sodium ascorbic acid, 2 MgSO4, 1 CaCl2, and 1 MgCl2. A mid-sagittal cut was made to divide the two hemispheres, and 300-μm coronal brain slices were cut using a vibrating blade microtome (Leica VT1200; Wetzlar, DE). The brain slices were transferred to a holding chamber with oxygenated aCSF containing (in mM): 119 NaCl, 2.5 KCl, 1 NaH2PO4, 26.2 NaHCO3, 11 d-glucose, 1 sodium ascorbic acid, 1.3 MgSO4, and 2.5 CaCl2 (∼295 mOsm, pH 7.2-7.3) at 37°C for 20 minutes and then room temperature for at least 40 minutes of rest for a total of one hour. The slices were kept submerged in oxygenated aCSF in a holding chamber at room temperature for up to 7-8 hours following slicing.

#### Electrophysiological recordings

All electrophysiological procedures were performed as previously described (Maria-Rios et al., 2023). After allowing the slices to rest for at least 1 h, individual slices were transferred to a recording chamber perfused with oxygenated aCSF (32 °C) containing 100 μM of the GABA_A_ receptor antagonist picrotoxin and 5 mM kynurenic acid to block glutamatergic transmission. Core and medial shell recordings from the NAc were made in the same slices obtained between +1.00 mm to +1.70 mm anterior from bregma (Paxinos and Franklin, 2019). The core and shell were distinguished based on their anatomical boundaries: the NAc core was identified as the region adjacent to the anterior commissure, while the medial shell was located ventromedial to the core, delineated by differences in cytoarchitecture and proximity to the lateral septum. Infrared differential interference contrast (IR-DIC) optics (Microscope: Olympus BX51; Camera: Dage-MIT) were used to visualize cells. Whole-cell current clamp recordings were performed using borosilicate glass pipettes (O.D. 1.5 mm, I.D. 0.86 mm; Sutter Instruments) with a 4-7-MΩ open tip resistance filled with a potassium gluconate-based internal solution containing (in mM): 122 K-gluconate, 20 HEPES, 0.4 EGTA, 2.8 NaCl, and 2 Mg^2+^ATP/0.3 Na_2_GTP (∼280 mOsm, pH adjusted to 7.2 with KOH). MSNs were identified based on morphology (medium-sized soma) as well as a hyperpolarized resting potential between -70 to -90 mV and inward rectification. Patched neurons were excluded based on a resting potential out of the desired range, characteristics of fast-spiking interneurons, and irregular firing patterns. All recordings were obtained using the MultiClamp 700B (Molecular Devices, San Jose, CA) amplifier and Digidata 1550A (Molecular Devices, San Jose, CA) digitizer. Data were filtered at 2 kHz, digitized at 10 kHz, and collected and analyzed using pClamp 10.0 software (Molecular Devices, San Jose, CA).

For whole-cell recordings, a membrane seal with a resistance >1 GΩ was achieved prior to breaking into the cell. Membrane capacitance (C_m_) and series resistance (R_s_) were recorded 1 minute after breaking in and were compensated under voltage-clamp. R_s_ was recorded in voltage-clamp with an average of 28 ± 6 MΩ upon entry and 26 ± 6 MΩ once the recording was finished for STs, 28 ± 7 MΩ and 27 ± 7 MΩ for GTs, and 30 ± 7 MΩ and 26 ± 7 MΩ for IRs (mean ± SD). Firing properties were recorded under current-clamp, and input resistance (R_i_) was monitored online during each sweep with a -100-pA, 25-ms current injection separated by 100 ms from the current injection step protocols. The average R_i_ across all sweeps is reported, and only cells with an R_i_ that remained stable (Δ < 20%) were included in the analysis (STs: n = 42, GTs: n = 40, IRs: n = 46). To assess firing properties, neurons were subjected to two recording protocols from their resting membrane potential (RMP). For spike number, spike frequency, voltage/current relationships, and sag ratios, neurons were subjected once to a step protocol consisting of 500-ms current injections starting from -500 pA to +500 pA in 25-pA increments with each sweep separated by 4 seconds. The RMP was calculated as the average voltage across all sweeps at 5 ms. Spike number was determined by counting individual action potentials at each level of current injection. Firing frequency (Hz) was calculated by averaging the frequency between consecutive spikes for a given current injection. If a neuron exhibited depolarization block, data for that cell were included up to the last current injection before the block occurred. Steady-state voltage responses were measured 200 ms after stimulus onset for each subthreshold current injection step. Sag ratios were calculated as the ratio of the peak voltage at the most hyperpolarized current injection (-500 pA) to the steady-state response. A ratio of 1.0 indicated no sag, with larger ratios corresponding to greater sag. The voltage/current relationship was assessed by measuring the difference between the steady-state voltage and the baseline voltage recorded 1 ms before stimulus onset.

To examine single action potential (AP) properties, neurons received 5-ms current injections in 25-pA increments, with each sweep separated by 4 seconds, until a single AP was elicited. The minimal current required to evoke an AP was defined as the current threshold (pA). AP threshold (mV) was determined as the voltage at the AP inflection point relative to 0 mV. The difference between RMP and AP threshold (Δ RMP/AP threshold, mV) was calculated by subtracting the AP threshold from the RMP. AP amplitude (mV) was measured as the voltage at the peak of the AP overshoot relative to 0 mV, and the difference between RMP and AP amplitude (Δ RMP/AP amplitude, mV) was also calculated. Similarly, the difference between AP threshold and AP amplitude (Δ AP threshold/AP amplitude, mV) was determined. AP half-width (ms) was defined as the duration of the AP at half of its peak amplitude.

## Experimental Design and Statistical Analysis

### Behavioral Studies

The behavioral response in the Pavlovian conditioned approach (PavCA) procedure was scored using an index that integrates the number, latency, and probability of lever presses (sign-tracking conditioned response) and magazine entries (goal-tracking conditioned response) during CS presentations within a session (Meyer et al., 2012). In brief, we averaged the response bias (i.e., number of lever presses and magazine entries for a session; [lever presses – magazine entries] / [lever presses + magazine entries]), latency score (i.e., average latency to perform a lever press or magazine entry during a session; [magazine entry latency – lever press latency]/10), and probability difference (i.e., proportion of lever presses or magazine entries; lever press probability – magazine entry probability). The index score ranges from +1.0 (absolute sign-tracking) to -1.0 (absolute goal-tracking), with 0 representing no bias. The average PavCA index scores of Sessions 5-6 were used to classify rats as STs (score ≥ 0.5), GTs (score ≤ -0.5), and IRs (-0.5 < score < 0.5). Out of a total of 31 rats, 10 were classified as STs, 7 were GTs, and 14 were IRs (**Figure 2B-D**).

GraphPad Prism 8 (Dotmatics) was used for statistical analysis of behavioral data. Number, latency, and probability for lever presses and for magazine entries, as well as PavCA index scores were analyzed using mixed-effects model via restricted maximum likelihood (REML). Fixed effects were set for phenotype (STs, GTs, IRs), session (1-6), and phenotype x session. Multiple comparisons were made using Sidak’s post-hoc test. Significance level was set at p < 0.05.

#### Electrophysiology Studies

A total of 128 cells from 31 rats were included in the analysis. The groups were divided as follows: STs group had a total of 10 rats, from which 23 cells were recorded in the core of 10 rats (1-2 slices/1-5 cells per rat) and 19 cells in the shell of 9 of the rats (1-2 slices/1-3 cells per rat); GTs group had a total of 7 rats, from which 17 cells were recorded in the core of 7 rats (1-2 slices/2-4 cells per rat) and 23 cells in the shell of 7 rats (1-2 slices/2-5 cells per rat); IRs group had a total of 14 rats, from which 31 cells were recorded in the core of 14 rats (1-2 slices/1-5 cells per rat) and 15 cells in the shell of 10 of the rats (1 slice/1-3 cells per rat).

For the analyses of sag ratio and voltage/current relationship curve, a total of 8 cells from STs (Core: n = 4, Shell: n = 4) and 17 cells from IRs (Core: n = 13, Shell: n = 4) were excluded. These cells underwent a step protocol ranging from -200 pA to +500 pA instead of the standard -500 pA to +500 pA, and were removed to maintain consistency in the hyperpolarized current injection analysis. Additionally, the single action potential protocol was not performed for 3 ST core cells, 1 GT core cell, 4 IR core cells, and 2 IR shell cells.

Electrophysiological data were analyzed offline using Clampfit 10.7 (Molecular Devices). Statistical analyses were conducted using GraphPad Prism 8 (Dotmatics) and SPSS Statistics (IBM). RMP, C_m_, R_i_, sag ratio, current to threshold, AP threshold, ΔRMP/AP threshold, AP amplitude, ΔRMP/AP amplitude, ΔAP threshold/AP amplitude, and AP halfwidth were analyzed separately for the core and shell using one-way ANOVA to assess group differences between STs, GTs, and IRs followed by Tukey’s post-hoc test for multiple comparisons (significance threshold: p < 0.05).

Number of spikes, spike firing frequency, and voltage/current (V/I) relationship curves (-500 to 0 pA and 0 to +100 pA) were analyzed using a linear mixed-effects model (REML), with significance set at p < 0.05. To assess phenotype differences within each subregion, fixed effects were defined for phenotype (STs, GTs, IRs), current injection (V/I: -500 pA to 0 pA, 0 pA to +100 pA; AP: +25 to +500 pA), and phenotype x current injection. Planned comparisons using mixed-effects model (REML) were conducted based on Sidak’s post-hoc test to obtain statistical values (F/P) for STs vs GTs, STs vs IRs, and GTs vs IRs.

For subregional analysis, a mixed-effects model (REML) was used with fixed effects set for phenotype (STs, GTs, IRs), subregion (core, shell), current injection (V/I: -500 pA to 0 pA, 0 pA to +100 pA; AP: +25 to +500 pA), subregion x phenotype, phenotype x current injection, subregion x current injection, and phenotype x subregion x current injection. Multiple comparisons were performed using Sidak’s post-hoc test. Based on the phenotype x subregion interaction, planned comparisons using a mixed-effects model (REML) were conducted to obtain subregion-specific statistics for STs (core vs shell), GTs (core vs shell), and IRs (core vs shell). Fixed effects for these comparisons included subregion (core, shell), current injection (V/I: -500 pA to 0 pA, 0 pA to +100 pA), and subregion x current injection.

For subregional differences within phenotypes in number of spikes and firing frequency, statistical values (F/P) for STs (core vs shell), GTs (core vs shell), and IRs (core vs shell), were derived from the phenotype x subregion Sidak’s post-hoc test.

## Results

### Following a PavCA procedure, rats were classified as STs, GTs, or IRs based on their lever and magazine response bias

To identify rats with ST, GT, and IR behavioral phenotypes in an outbred population, all rats underwent six days of PavCA training with a lever-cue (CS) paired with the response-independent delivery of a banana-flavored food pellet into a magazine (US) (**Figure 2A-C**). Based on their PavCA index score on Sessions 5–6, rats were classified as either STs (score ≥ 0.5; n = 10), GTs (score ≤ -0.5; n = 7), or intermediate responders (-0.5 < score < 0.5; n = 14) (**Figure 2D**). A significant session × phenotype interaction was observed for the PavCA index score (F(10,138) = 19.0, p < 0.0001), indicating a strong difference in behavioral response bias toward the lever or magazine that develops across training sessions, suggesting differences in the attribution of incentive and predictive properties to the cue (**Figure 2E**). Post hoc comparisons confirmed that all groups significantly differed from one another (p < 0.0001). A significant session × phenotype interaction was found in lever press number (F(10,138) = 12.9, p < 0.0001), latency (F(10,138) = 12.4, p < 0.0001), and probability (F(10,138) = 8.06, p < 0.0001), indicating that the rate of acquisition differed among phenotypes (**Figure 3A-C**). Post hoc comparisons revealed that STs pressed the lever significantly more than GTs (p < 0.0001) and IRs (p < 0.0001), while IRs exhibited more lever pressing than GTs (p < 0.0001). This pattern aligns with previous findings demonstrating that STs readily approach and interact with reward-predictive cues, whereas GTs direct their behavior toward the location of reward delivery (Flagel et al., 2009; Maria-Rios et al., 2019). The inclusion of multiple behavioral parameters allows for a more comprehensive characterization of response acquisition, highlighting the distinct learning strategies employed by each phenotype. Conversely, a significant session × phenotype interaction was also observed for magazine entry responses (F(10,138) = 13.7, p < 0.0001), latency (F(10,138) = 15.1, p < 0.0001), and probability (F(10,138) = 20.3, p < 0.0001). Post hoc comparisons revealed that STs exhibited significantly fewer magazine entries than GTs (p = 0.0041) and IRs (p = 0.015), while IRs and GTs did not differ from each other (p = 0.54) (**Figure 3D-F**). This finding is consistent with historical data from our lab and others, reinforcing that GTs rely more on goal-directed behavior, attending to the reward location rather than the cue itself (Flagel et al., 2009; Maria-Rios et al., 2019).

**Figure 3.**
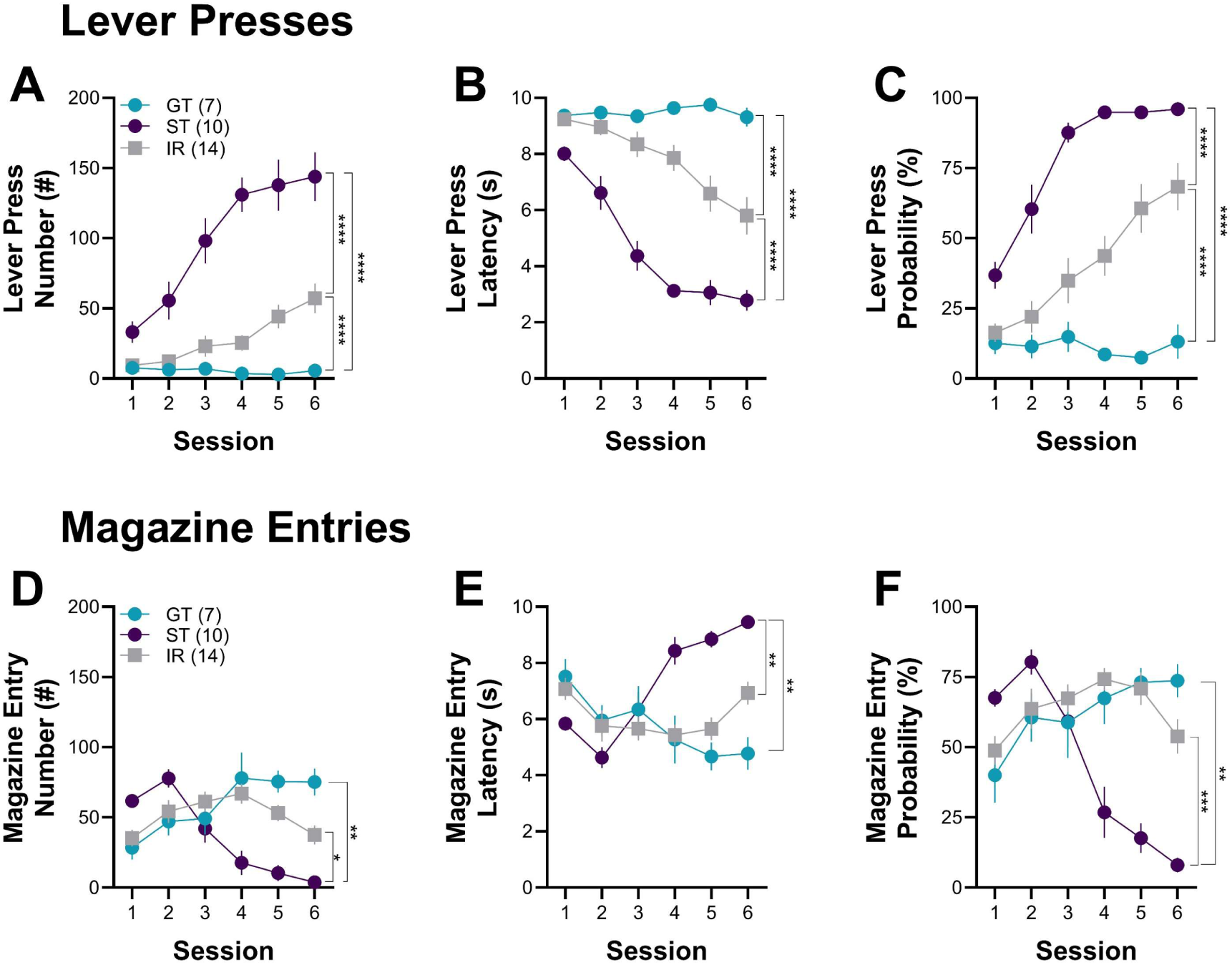
Behavioral differences in lever pressing and magazine entries during PavCA. STs and GTs differed in the number, latency, and probability of lever presses and magazine entries during six PavCA sessions. (**A**) Lever press number, (**B**) latency, and (**C**) probability over sessions for STs, GTs, and IRs. (**D**) Magazine entry number, (**E**) latency, and (**F**) probability across sessions. STs showed higher lever press number, lower latency, and greater probability compared to both GTs and IRs. GTs and IRs displayed greater magazine entries, lower latency, and higher probability than STs, with no difference between GTs and IRs. Significance for Sidak’s post hoc test is shown as *p < 0.05, **p < 0.01, ***p < 0.001, ****p < 0.0001. Data are presented as mean ± S.E.M.

### Passive membrane properties of NAc MSNs in the core and shell subregions of STs, GTs, and IRs rats

To assess whether STs, GTs, and IRs exhibit differences in intrinsic excitability, we examined the passive membrane properties of MSNs in the NAc core and shell (**Figure 4A**). Whole-cell patch-clamp recordings were performed in rat brain slices following a 1–3 week resting period from the last PavCA training session (**Figure 1**). This resting period was intended to allow training-induced neuronal changes to return to baseline so that our measurements would reflect innate differences in intrinsic excitability across the three phenotypes. No significant main effects of phenotype were observed in the NAc core for resting membrane potential (RMP) (F(2,68) = 0.55, p = 0.58) or input resistance (F(2,68) = 0.33, p = 0.72) (**Figures 4B–C**). However, in the NAc shell, a significant main effect of phenotype was detected for RMP (F(2,54) = 4.98, p = 0.010). Post hoc analysis indicated that STs exhibited a significantly more hyperpolarized RMP compared to GTs (p = 0.0079), while IRs did not differ significantly from the other groups (**Figure 4D**). Input resistance remained non-significant across phenotypes (F(2,54) = 1.96, p = 0.15) (**Figure 4E**). These findings suggest that intrinsic excitability differences between phenotypes may be more prominent in the NAc shell than in the core, with STs exhibiting a more hyperpolarized RMP. This could indicate a lower baseline excitability in ST MSNs, potentially contributing to differences in how reward-related cues are processed and attributed with incentive salience.

**Figure 4.**
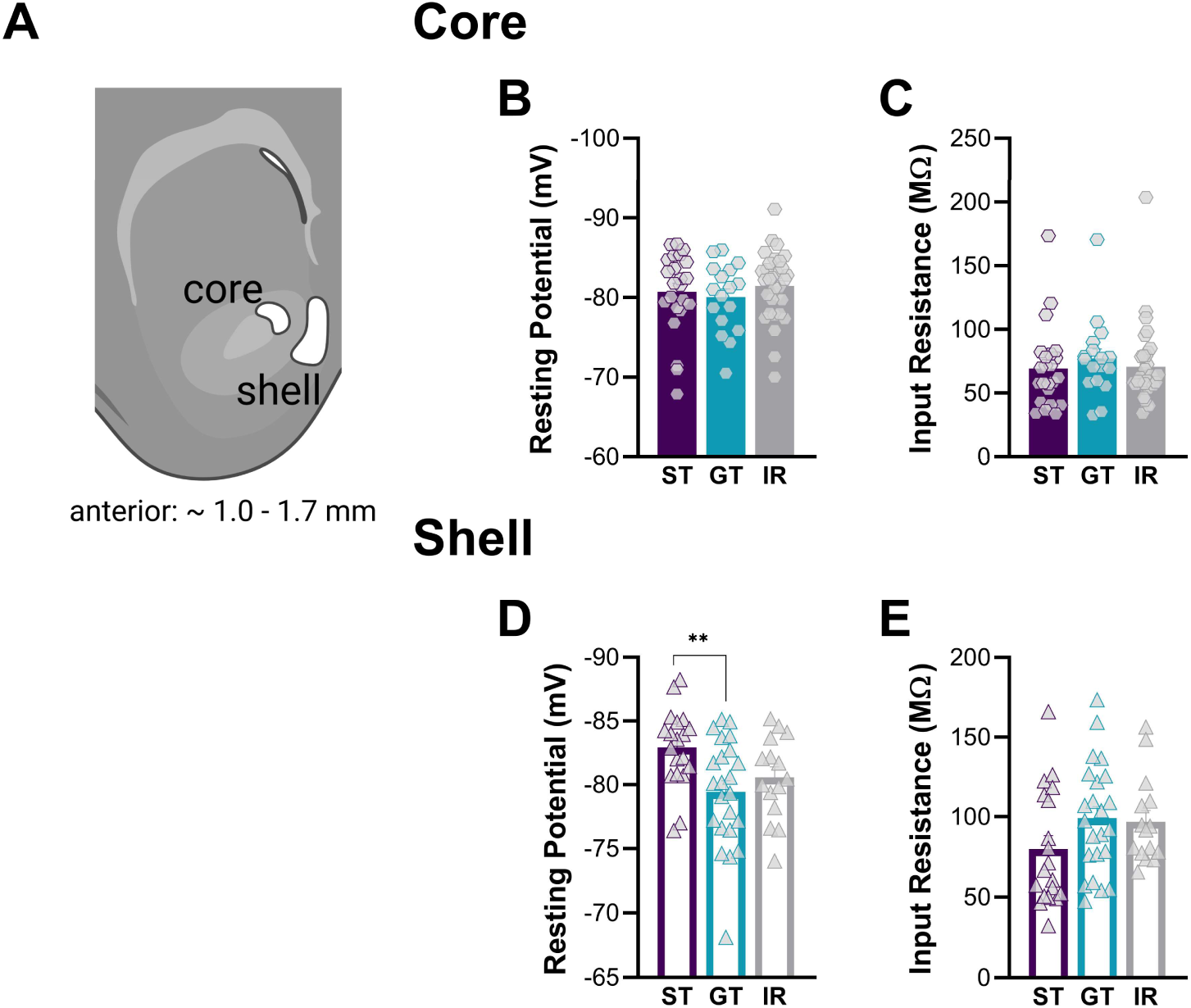
Membrane properties of NAc MSNs in ST, GT, and IR rats. (**A**) Diagram showing the NAc core and shell subregions where whole-cell recordings were made. (**B**) Resting potential and (**C**) input resistance in the NAc core showed no significant differences between STs, GTs, and IRs. (**D**) In the NAc shell, STs had a more hyperpolarized resting membrane potential compared to GTs. (**E**) No differences in input resistance were observed across groups in the NAc shell. Significance for Tukey’s post hoc test is shown as **p < 0.01. Data are presented as mean ± S.E.M.

### NAc MSNs in the core and shell of STs exhibit lower hyperpolarization than GTs and IRs in response to negative current inputs

We next examined whether active intrinsic excitability properties differed across phenotypes. To assess this, MSNs in the NAc core and shell were subjected to a current injection step protocol ranging from -500 to +500 pA in 25-pA increments (**Figure 5A, 5D**). We first analyzed the voltage/current (V/I) relationship for hyperpolarizing currents (-500 to 0 pA) to determine whether the neurons’ responses to negative current injections varied by phenotype. A significant main effect of phenotype was found in the V/I relationship for hyperpolarizing currents (-500 to 0 pA) in both the NAc core (F(2,1134) = 13.4, p < 0.0001) and shell (F(2,966) = 20.9, p < 0.0001) (**Figures 5B, 5E**), indicating that MSNs from different phenotypes exhibit distinct passive responses to negative current inputs. Post hoc comparisons revealed that ST MSNs exhibited significantly lower hyperpolarization in response to negative current injections compared to GTs (p < 0.0001) and IRs (p < 0.0001) in both regions. This difference in hyperpolarization response could influence how reward-related information is processed in the NAc. In contrast, when examining responses to depolarizing current injections (0 to 100 pA), no significant main effect of phenotype was found in either the core (F(2,265) = 1.60, p = 0.20) or shell (F(2,221) = 2.81, p = 0.063), suggesting that responses to positive currents were comparable across phenotypes. This suggests that while ST MSNs differ in their hyperpolarization response, their response to positive current remain largely unchanged relative to GTs and IRs.

**Figure 5.**
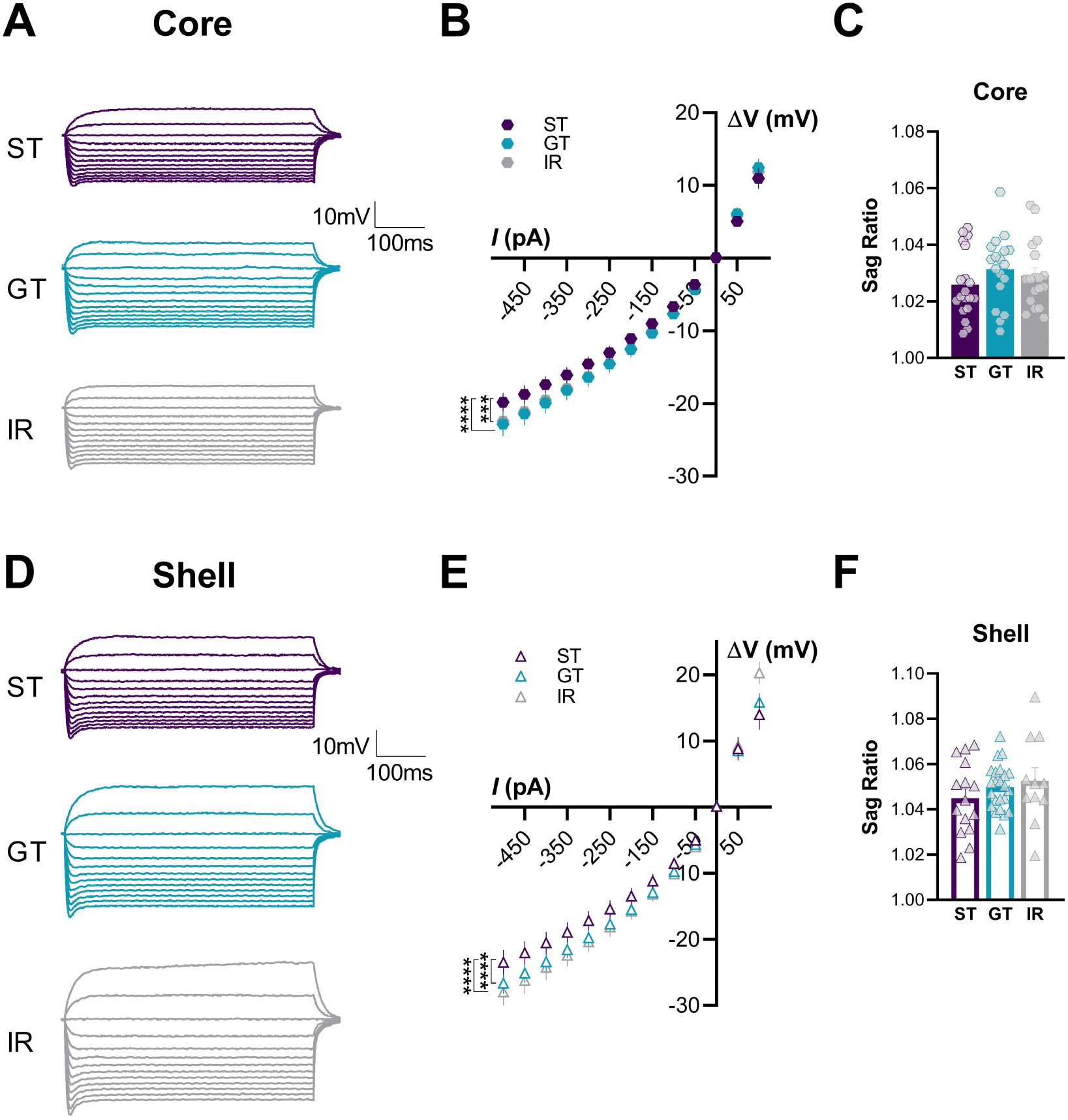
Differential hyperpolarization responses of NAc MSNs in STs, GTs, and IR. (**A, D**) Representative voltage traces from current-clamp recordings of MSNs in the NAc core and shell from STs, GTs, and IRs. (**B, E**) Voltage/current relationship: STs exhibited significantly lower hyperpolarization in response to negative current injections compared to GTs and IRs in both the NAc core and shell. No differences were found between phenotypes in response to depolarizing current injections. (**C, F**) Sag ratio: No significant differences were observed between phenotypes in the sag ratio in the NAc core or shell. Significance for mixed-effects model planned comparisons is shown as *** p < 0.001, **** p < 0.0001. Data are presented as mean ± S.E.M.

To further examine differences in neuronal excitability, we analyzed the sag response, a measure of how neurons recover from hyperpolarization and regulate their membrane potential following inhibitory input (Pape, 1996; Robinson and Siegelbaum, 2003). The sag ratio was obtained by dividing the peak voltage response to -500 pA over the steady-state response 200 ms from stimulus onset. No significant differences were observed across phenotypes (Core: F(2,52) = 1.01, p = 0.37; Shell: F(2,46) = 0.97, p = 0.39) (**Figures 5C, 5F**). This suggests that MSNs across phenotypes exhibit similar capacities to stabilize their membrane potential after hyperpolarizing inputs.

Overall, across all STs, GTs, and IRs, NAc MSNs in the shell exhibited significantly greater excitability than core MSNs, as both hyperpolarizing and depolarizing current injections consistently induced a larger change in membrane potential in the shell (Hyperpolarizing: F(1,2100) = 175, p < 0.0001; Depolarizing: F(1,486) = 59.7, p < 0.0001). This aligns with prior research demonstrating that the NAc shell is generally more excitable than the core (Kourrich et al., 2007; Maria-Rios et al., 2023), possibly reflecting differences in synaptic input integration and output connectivity between these subregions.

### NAc MSNs in the core and shell of STs, GTs, and IRs exhibit distinct firing properties

To further characterize the intrinsic excitability of MSNs from the core and shell of STs, GTs, and IRs, we examined their firing properties in response to current injection steps ranging from +25 to +500 pA in 25-pA increments (**Figure 6A, 6D**). A significant main effect of phenotype was observed for the number of spikes in both the NAc core (F(2,1316) = 19.3, p < 0.0001) and shell (F(2,1058) = 12.1, p < 0.0001) (**Figures 6B, 6E**), suggesting that MSNs from different phenotypes exhibit distinct excitability profiles. Post hoc comparisons revealed phenotype-specific differences in excitability. In the core, GT MSNs exhibited significantly more spikes than both STs (p < 0.0001) and IRs (p < 0.01), while STs had the lowest excitability (**Figure 6B**). In contrast, in the NAc shell, IR MSNs fired significantly more spikes than both STs (p < 0.0001) and GTs (p < 0.001), but STs and GTs did not differ from one another (**Figure 6E**). A similar pattern was observed for firing frequency (Core: F(2,1316) = 20.4, p < 0.0001; Shell: F(2,1058) = 12.1, p < 0.0001) (**Figures 6C, 6F**). These findings indicate that neuronal excitability varies across phenotypes in a region-dependent manner, although the behavioral implications of these differences remain to be further explored.

**Figure 6.**
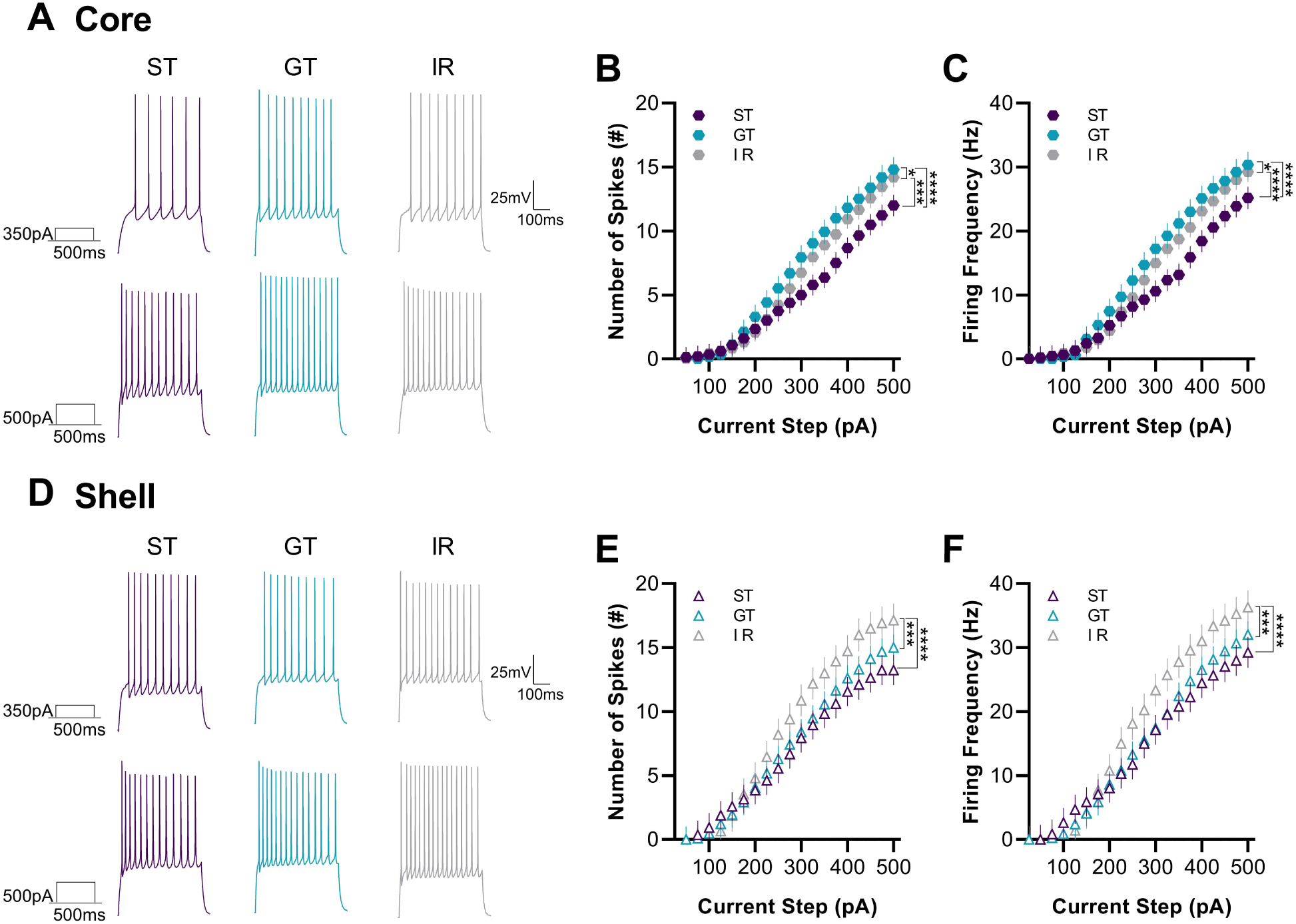
Distinct firing properties of NAc MSNs in STs, GTs, and IRs. (**A, D**) Representative current-clamp traces from MSNs in the NAc core and shell following 350-pA (top) and 500-pA (bottom) current injections in STs, GTs, and IRs. (**B, C**) In the NAc core, GTs exhibited the highest number of spikes and firing frequency, while STs had the lowest, with significant differences between phenotypes. (**E, F**) In the NAc shell, IRs had significantly higher spikes and firing frequency compared to both STs and GTs, with no differences between STs and GTs. Significance for mixed-effects model planned comparisons is shown as *p < 0.05, ***p < 0.001, ****p < 0.0001. Data are presented as mean ± S.E.M.

We also found significant subregional differences in firing properties between the core and shell. A significant phenotype × subregion interaction was found for number of spikes (F(2,2374) = 10.8, p < 0.0001), indicating that subregional differences in excitability varied across phenotypes. Post hoc comparisons confirmed that there was greater firing in the shell than in the core in STs (p < 0.0001) and IRs (p < 0.0001), but not in GTs (p = 0.078). While the functional significance of these differences remains unclear, they may reflect variations in how each phenotype integrates and responds to motivationally relevant cues.

Lastly, we examined whether specific properties of the action potentials (APs) of MSNs in the core and shell were different between STs, GTs, and IRs. We delivered 5-ms current injections in 25-pA increments until a single AP was elicited and found that the action potential properties in both subregions did not differ between STs and GTs (data not shown in figure; see **Table 1** for a full list of AP properties). In summary, no differences were observed in the current required to elicit a single AP, AP halfwidth, AP threshold from zero and resting potential, or AP peak amplitude from zero, resting potential, and threshold (p > 0.05). However, we identified subtle differences between STs and IRs. In the core, STs exhibited a significantly longer AP halfwidth than IRs (p = 0.015), suggesting slower repolarization. In the shell, IRs had a more depolarized AP threshold (p = 0.03), and a consequently smaller AP peak amplitude measured from threshold (p = 0.027).

**Table 1.** Electrophysiological passive and active properties of medium spiny neurons in the core and shell of nucleus accumbens of STs, GTs, and IRs rats. Table lists mean ± S.E.M. (sample size) for passive and active properties of core and shell MSNs for STs, GTs, and IRs. Significance for Tukey’s post hoc test between STs and IRs is shown as *p < 0.05.

|  | Core |  |  | Shell |  |  |
| --- | --- | --- | --- | --- | --- | --- |
|  | Sign-trackers | Goal-trackers | Intermediates | Sign-trackers | Goal-trackers | Intermediates |
| *Passive membrane properties* |  |  |  |  |  |  |
| Resting membrane potential, mV | -81 ± 1 (23) | -80 ± 1 (17) | -81.5 ± 0.8 (31) | -82.9 ± 0.7 (19) | -79.4 ± 0.9 (23) | -80.6 ± 0.8 (15) |
| Cell capacitance, pF | 150 ± 8 (23) | 130 ± 7 (17) | 122 ± 5 (31) | 105 ± 6 (19) | 88 ± 4 (23) | 86 ± 6 (15) |
| Input resistance, MΩ | 69 ± 7 (23) | 77 ± 7 (17) | 71 ± 6 (31) | 80 ± 8 (19) | 99 ± 7 (23) | 97 ± 7 (15) |
| *Active membrane properties* |  |  |  |  |  |  |
| V/I curve (-500pA to 0pA) | -11.9 ± 0.2 (19) | -13.6 ± 0.2 (18) | -13.3 ± 0.2 (18) | -14.3 ± 0.3 (15) | -16.4 ± 0.2 (23) | -16.9 ± 0.3 (11) |
| V/I curve (0pA to +100pA) | 5.3 ± 0.4 (19) | 6.2 ± 0.4 (18) | 5.6 ± 0.4 (18) | 7.7 ± 0.6 (15) | 8.4 ± 0.5 (23) | 9.8 ± 0.6 (11) |
| Sag ratio at -500pA, mV | 1.026 ± 0.003 (19) | 1.031 ± 0.003 (18) | 1.029 ± 0.003 (18) | 1.045 ± 0.004 (15) | 1.050 ± 0.002 (23) | 1.053 ± 0.006 (11) |
| Number of spikes, AP# | 4.7 ± 0.2 (23) | 6.4 ± 0.2 (17) | 5.7 ± 0.2 (31) | 6.5 ± 0.3 (19) | 7.0 ± 0.2 (23) | 8.3 ± 0.3 (15) |
| Firing frequency, Hz | 10.0 ± 0.4 (23) | 13.7 ± 0.4 (17) | 12.2 ± 0.3 (31) | 14.0 ± 0.5 (19) | 14.6 ± 0.5 (23) | 17.7 ± 0.6 (15) |
| Current to threshold, pA | 981 ± 56 (20) | 875 ± 60 (16) | 879 ± 42 (27) | 836 ± 63 (19) | 711 ± 45 (23) | 694 ± 40 (13) |
| AP threshold, mV | -45 ± 1 (20) | -44 ± 1 (16) | -45 ± 1 (27) | -45 ± 2 (19)* | -41 ± 1 (23) | -39 ± 2 (13) |
| $\Delta$ RMP/AP threshold, mV | 36 ± 2 (20) | 35 ± 1 (16) | 37 ± 8 (27) | 37 ± 2 (19) | 38 ± 1 (23) | 42 ± 2 (13) |
| AP amplitude, mV | 53.9 ± 0.9 (20) | 53 ± 1 (16) | 51 ± 1 (27) | 53 ± 1 (19) | 53 ± 2 (23) | 49 ± 2 (13) |
| $\Delta$ RMP/AP amplitude, mV | 135 ± 1 (20) | 132 ± 2 (16) | 132 ± 1 (27) | 135 ± 1 (19) | 133 ± 2 (23) | 130 ± 2 (13) |
| $\Delta$ AP threshold/AP amplitude, mV | 99 ± 2 (20) | 97 ± 2 (16) | 96 ± 2 (27) | 97 ± 2 (19)* | 94 ± 2 (23) | 88 ± 3 (13) |
| AP halfwidth, ms | 0.72 ± 0.02 (20)* | 0.68 ± 0.02 (16) | 0.65 ± 0.01 (27) | 0.70 ± 0.02 (19) | 0.68 ± 0.02 (23) | 0.69 ± 0.02 (13) |

These findings highlight distinct electrophysiological properties of MSNs in the NAc core and shell across STs, GTs, and IRs, reflecting phenotype-specific differences in intrinsic excitability and firing patterns.

## Discussion

This is the first study to demonstrate that MSNs in the NAc of STs, GTs, and IRs exhibit distinct intrinsic membrane properties, with differences observed across the core and shell subregions. Most notably, STs had the lowest intrinsic excitability in the NAc core, while GTs exhibited the highest. Although STs and GTs did not differ in the shell firing properties, IRs showed greater intrinsic excitability than both phenotypes. Given that the NAc core encodes the motivational value of conditioned cues, while the shell modulates unconditioned responses (Meredith et al., 2008; Saunders et al., 2018), it is unsurprising that the strongest phenotype differences emerged in the core.

These findings suggest that the previously reported differences in NAc core and shell activity during Pavlovian learning may be linked to the intrinsic excitability state of their MSNs. STs show increased cue-evoked activity in both the NAc core and shell (Flagel et al., 2011a), alongside a reduction in reward-evoked activity over PavCA training (Gillis and Morrison, 2019). This pattern, often interpreted as a prediction error signal, is notably absent in GTs despite their clear learning of cue-reward relationship**s**. This has led to the hypothesis that the shift from reward-evoked to cue-evoked activity in the NAc may primarily reflect the attribution of incentive properties to a reward-paired cue as opposed to predictive properties. Here, we found that GTs exhibited the highest intrinsic excitability in the NAc core, raising the possibility that a higher neuronal excitability state in GTs may make it more difficult for their incentive salience-related NAc core activity to shift away from the reward and toward the cue. Thus, some of the differences in reward learning processes between GTs and STs might be traced to subregional differences in the intrinsic neurophysiological properties of their NAc core neurons.

Reduced intrinsic excitability of MSNs is a well-established feature of cocaine exposure and withdrawal (for review see Wolf 2010). Lower MSN excitability has been directly linked to increased cocaine sensitization and self-administration (Dong et al., 2006; Mu et al., 2010). Behaviorally, STs exhibit greater addiction-like behaviors than GTs, including enhanced psychomotor sensitization to cocaine (Flagel et al., 2008), higher cocaine preference over food (Tunstall and Kearns, 2015), and increased cue-induced reinstatement of cocaine and nicotine (Saunders and Robinson, 2011; Versaggi et al., 2016). Therefore, reduced MSN excitability in STs may be a neurobiological feature that predisposes them to pathological drug responses.

The decrease in MSN excitability following cocaine exposure and withdrawal has been linked to reductions in voltage-dependent Na^+^ (Zhang et al., 1998) and Ca^2+^ (Zhang et al., 2002) currents, as well as increased K^+^ channel activity (Hu et al., 2004). One candidate mechanism is SK-type Ca^2+^- activated K^+^ channels, as blocking these channels with apamin partially prevents cocaine-induced reductions in MSN excitability (Ishikawa et al., 2009). Interestingly, inositol and corticosterone – neurochemicals that differ between STs and GTs (Flagel et al., 2009; Fitzpatrick et al., 2016) – can modulate SK channel activity, increasing K^+^ currents and reducing neuronal excitability (Yamada et al., 2004; Kye et al., 2007; Clements et al., 2013). This raises the possibility that the lower intrinsic excitability observed in STs, even in the absence of drug exposure, may be linked to differences in K+ channel regulation. Future studies should explore whether SK-type K+ channels contribute to phenotype-specific differences in MSN excitability.

It is important to note that most of the mentioned studies reporting a reduction in MSN excitability have been in the NAc shell. While STs and GTs differed in firing properties primarily in the core, ST MSNs in the shell exhibited a significantly more hyperpolarized resting membrane potential, suggesting a lower baseline excitability. Additionally, both STs and GTs showed reduced shell excitability compared to IRs. Few studies have examined IRs in behavioral and neurobiological contexts, making it difficult to predict how their excitability differences relate to behavior. It is known that appetitive Pavlovian conditioning can increase the intrinsic excitability of specific neuronal ensembles of MSNs in the NAc shell (Ziminski et al., 2017). It is possible that an overall greater intrinsic excitability in the NAc shell of IRs may facilitate the attribution of both predictive and incentive properties to cues, giving them more flexibility in their conditioned responses. However, additional studies are needed to characterize IRs at this level.

Dopaminergic transmission strongly influences MSN excitability in the NAc (O’Donnell and Grace, 1996; Nicola et al., 2000; Surmeier et al., 2007; Planert et al., 2013). During Pavlovian learning, STs and GTs exhibit distinct patterns of dopamine (DA) release, with DA responses mirroring cue- and reward-evoked activity in the NAc (Flagel et al., 2011b). Additionally, DA agonists and antagonists differentially affect STs and GTs (Saunders and Robinson, 2012; Chow et al., 2016), suggesting pre-existing differences in the dopaminergic system. Indeed, some studies have reported differences in the DA system of STs and GTs that may modulate the way MSNs respond to DA, and therefore affect their intrinsic membrane excitability properties (Flagel et al., 2007; Singer et al., 2016). For example, STs have greater dopamine transporter (DAT) expression in the NAc core than GTs (Singer et al., 2016), leading to faster DA clearance from the synapse, independent of baseline DA release levels (Singer et al., 2016). This rapid DA re-uptake is thought to be important for more time-locked responses of MSNs to phasic DA release during cue presentations and may help potentiate the value of incentive stimuli and attribution of incentive salience (Wieland et al., 2015; Singer et al., 2016). Interestingly, activation of D_1_-like receptors in MSNs can cause an increase in MSN depolarization by inhibiting Kir-channel K^+^ currents (Podda et al., 2010) and by enhancing L-type Ca^2+^ currents (Hernández-López et al., 1997). Therefore, if GTs have greater tonic DA levels in the NAc, this may lead to stronger D1 receptor activation increasing membrane excitability compared to STs. Lower intrinsic excitability in STs, combined with a more fine-tuned response to DA release may selectively enhance the response of MSNs to only the most salient stimuli and thereby promote the attribution of incentive salience to specific stimuli. Additionally, D_1_-mediated excitability occurs mainly when MSNs are already near threshold (up-state), which may explain why GT MSNs exhibit higher excitability in depolarized but not hyperpolarized states.

We found that subregional differences between core and shell MSNs were larger for STs and IRs than for GTs. In both STs and IRs, MSNs in the shell fired more action potentials than those in the core, whereas GTs showed no significant subregional difference. It is interesting that IRs share this feature with STs. In this case, this shared characteristic may suggest that a subregional difference might be important for sign-tracking responses. Future studies are needed to test whether experimental manipulation of subregional differences between core and shell excitability can affect individual differences in the attribution of incentive salience.

A key limitation of this study is the inability to determine whether MSN excitability differences in STs and GTs are innate or shaped by PavCA training. This is because currently, there is no method to distinguish STs from GTs before training, making it unclear whether observed excitability differences are pre-existing or experience-dependent. First, the training experience itself (e.g., handling, reward exposure) may be enough to alter NAc physiology (Scala et al., 2018) and second, the actual learning component (e.g., CS-US associations) of the task may also have a distinct impact (Ziminski et al., 2017). We attempted to control for these variables by waiting a period of at least one week before the electrophysiological recordings with the hopes that any training-induced changes would most likely have returned to baseline by then. Based on this limitation, although we found clear differences between STs, GTs, and IRs in membrane properties of MSNs, we could not determine whether these are pre-existent differences and conclude that all three phenotypes are innately different. To determine whether individual differences in PavCA behavior are innate and caused by intrinsic differences in NAc excitability, some method would be needed to identify STs and GTs prior to PavCA training, and/or measurement of MSN excitability would have to be done *in vivo* to test how excitability changes over the course of behavioral training.

The overwhelming evidence suggests that neuronal activity in the NAc is particularly important for individual differences in the attribution of incentive salience. It is very likely that the distinct patterns of activity during PavCA, are the collective result of differences in inputs to the NAc modulated by both internal and external stimuli in STs and GTs. However, our results suggest that these characteristic patterns of activity can be directly influenced by differences in the intrinsic neuronal properties of MSNs in the NAc of STs and GTs. This in turn may help understand how inputs to the NAc may exert different synaptic influence because of baseline differences in its excitability state. We intend for these studies to aid in our understanding on the neurobiology of individual differences in incentive salience attribution and endophenotypes associated with addiction vulnerability.

## Acknowledgments

This work was funded by the the National Science Foundation Graduate Research Fellowship Program (CEM), the National Institute of Neurological Disorders and Stroke (NINDS; F99 NS120544 [CEM]) and the National Institute on Drug Abuse (NIDA; K08 DA037912 [JDM]; R01 DA044960 [JDM]; T32 DA007281 [CJF]).

## Author Contributions

Author contributions: C.E.M., G.G.M., and J.D.M. designed research; C.E.M. performed research; C.E.M. analyzed data; C.E.M., G.G.M., and J.D.M. wrote the paper.

